## Supplemental figures 1-6 for "Nse5/6 inhibits the Smc5/6 ATPase to facilitate DNA substrate selection"

A

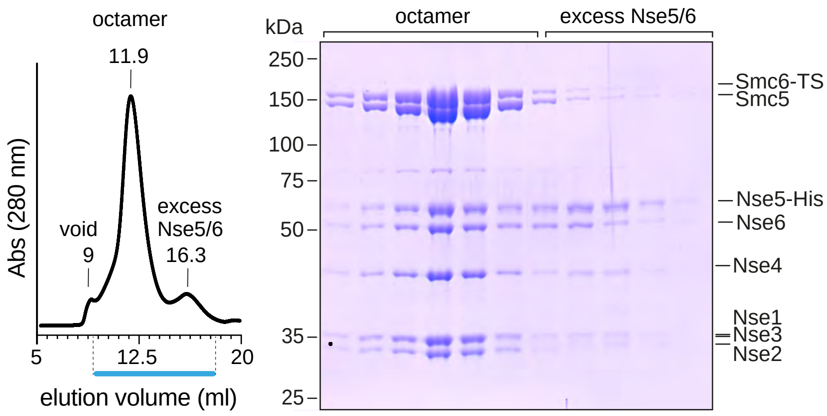


B

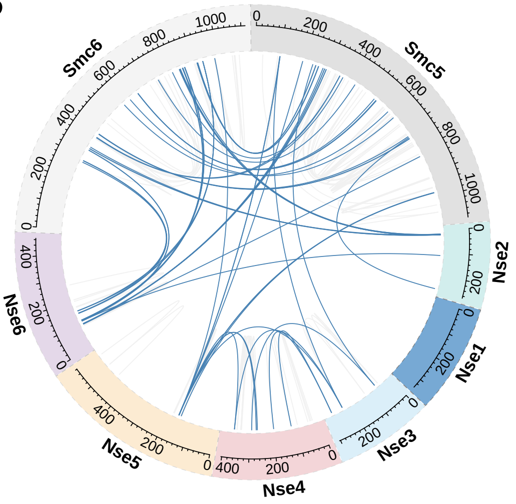


C

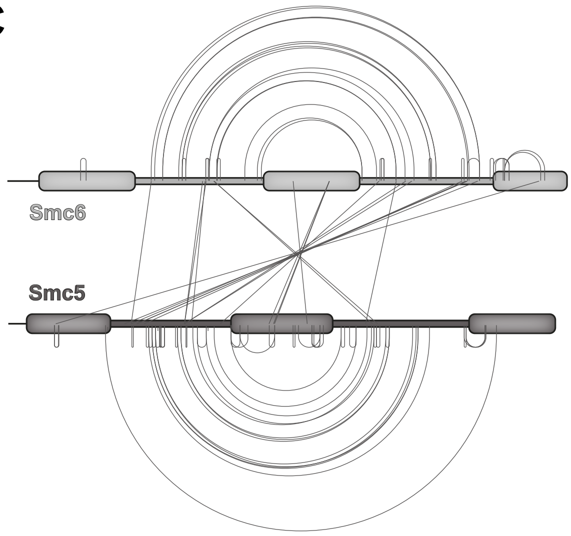


### SUPPLEMENTAL FIGURE 1

Fig S1 Reconstitution and architecture of Smc5/6

A. Gel filtration of Smc5/6 holo-complexes reconstituted by mixing Nse5/6 dimer and Smc5/6 hexamer roughly at a 3:2 ratio. Elution profiles measured by absorption at 280 nm (*left panel*). Selected fractions were analysed by SDS-Page and Coomassie staining (*right panel*).

B. Circular representation of lysine-lysine inter-subunits cross-links identified by mass spectrometry (XL-MS) in buffer containing 250 mM NaCl. As in Fig 1B but all inter- and intra-subunit cross-links are shown as individual lines. Intralinks are in light grey, interlinks in blue colours.

C. Cross-links in the Smc5(EQ)/Smc6(EQ) hexamer in the absence of substrates as identified by XL-MS. Display as in Fig 1C. Equivalent data for the wild-type Smc5/6 octamer is shown in Fig 1C.

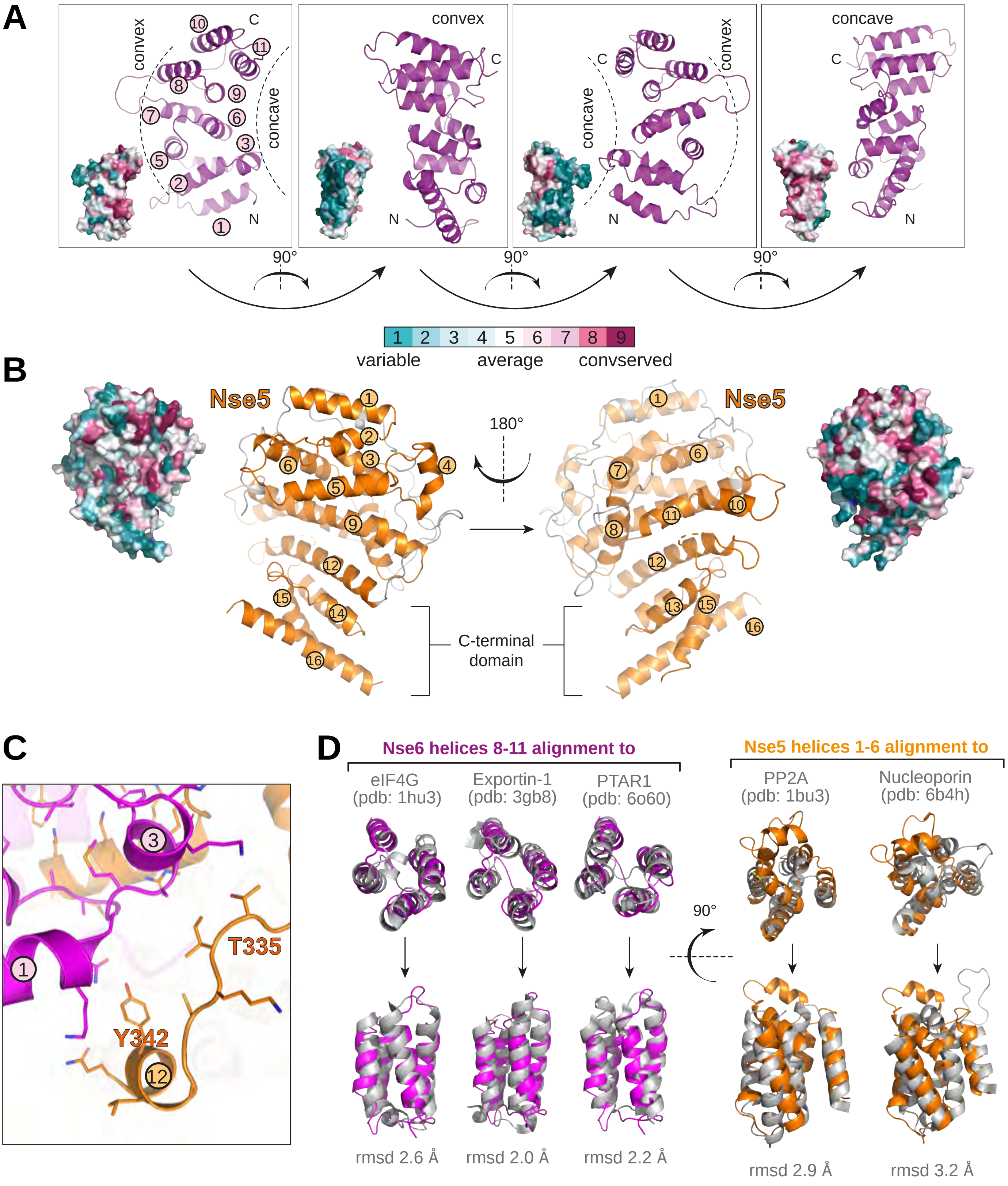


**SUPPLEMENTAL FIGURE 2**

Fig S2 Structural views and conservation of Nse5 and Nse6

A. The Nse6 moiety of the Nse5/6 dimer structure is shown in front, back, and side views in cartoon representation and in corresponding surface conservation displays (at the bottom left of each panel). The concave surface interacting with Nse5 shows highest residue conservation. Conservation colour code is indicated at the bottom.

B. Front and back views of Nse5 in the Nse5/6 structure in cartoon and corresponding surface conservation representation. Display as in (A). Conservation colour code as in (A).

C. Zoom view of contacts at the Nse5/6 interface formed by residues in helix α12 as well as the preceding loop in Nse5 with Nse6 helices α1 and α3.

D. Superimposition of Nse6 helices α8-α11 with selected top hits from a DALI search in the Protein Data Bank (*left panel*) in top and front view. Root mean square displacement (RMSD) values are given as indicator for the quality of the fit.

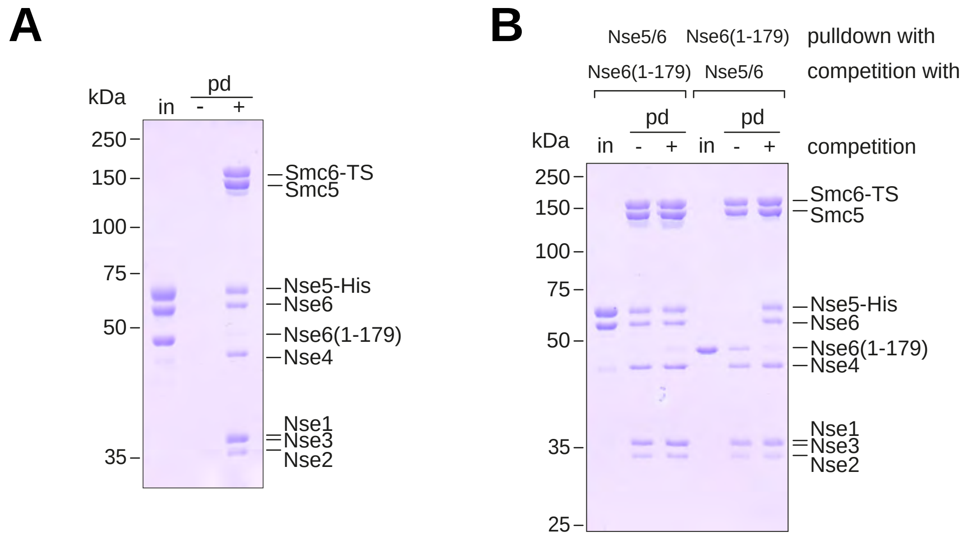


### SUPPLEMENTAL FIGURE 3

Fig S3 Smc5/6 interactions measured by pulldowns

A. Competition binding of a mixture of Nse5-His/Nse6 and Nse6(1-179)-CPD-His (‘in’) to immobilized Smc5/Smc6-Twin-Strep hexamer. Input (‘in’) and pulldown fractions were analysed by SDS-Page and Coomassie staining.

B. Competition with excess Nse5/6 and Nse6 fragment. Smc5/6 hexamers were immobilized together with Nse5-His/Nse6 or Nse6(1-179)-CPD-His and washed with Nse6(1-179)-CPD-His or Nse5-His/Nse6, respectively. Input (‘in’), pulldown and control pulldowns (without competition) were analysed by SDS-Page and Coomassie staining.

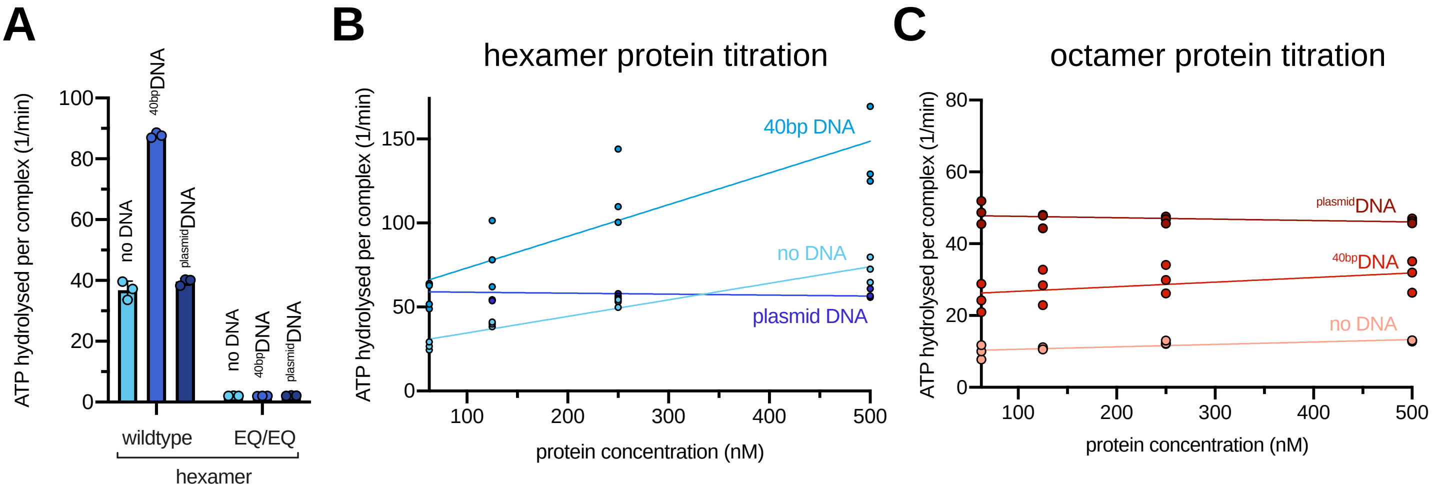


### SUPPLEMENTAL FIGURE 4

Fig S4 ATP hydrolysis by Smc5/6 and mutants thereof

A. ATP hydrolysis rates for wild-type Smc5/6 hexamers and the Smc5(EQ)/Smc6(EQ) variant. Rates for the absence and presence of DNA substrates are given (per Smc complex per minute).

B. Cooperativity in ATP hydrolysis by the Smc5/6 hexamer. ATP hydrolysis rates were determined at different protein concentrations. Smc5/6 hexamers showed cooperative behaviour without DNA and with ${}^{40bp}$DNA but not with ${}^{plasmid}$DNA.

C. Absence of cooperative in ATP hydrolysis by the Smc5/6 octamer. As in (B) using reconstituted octamers.

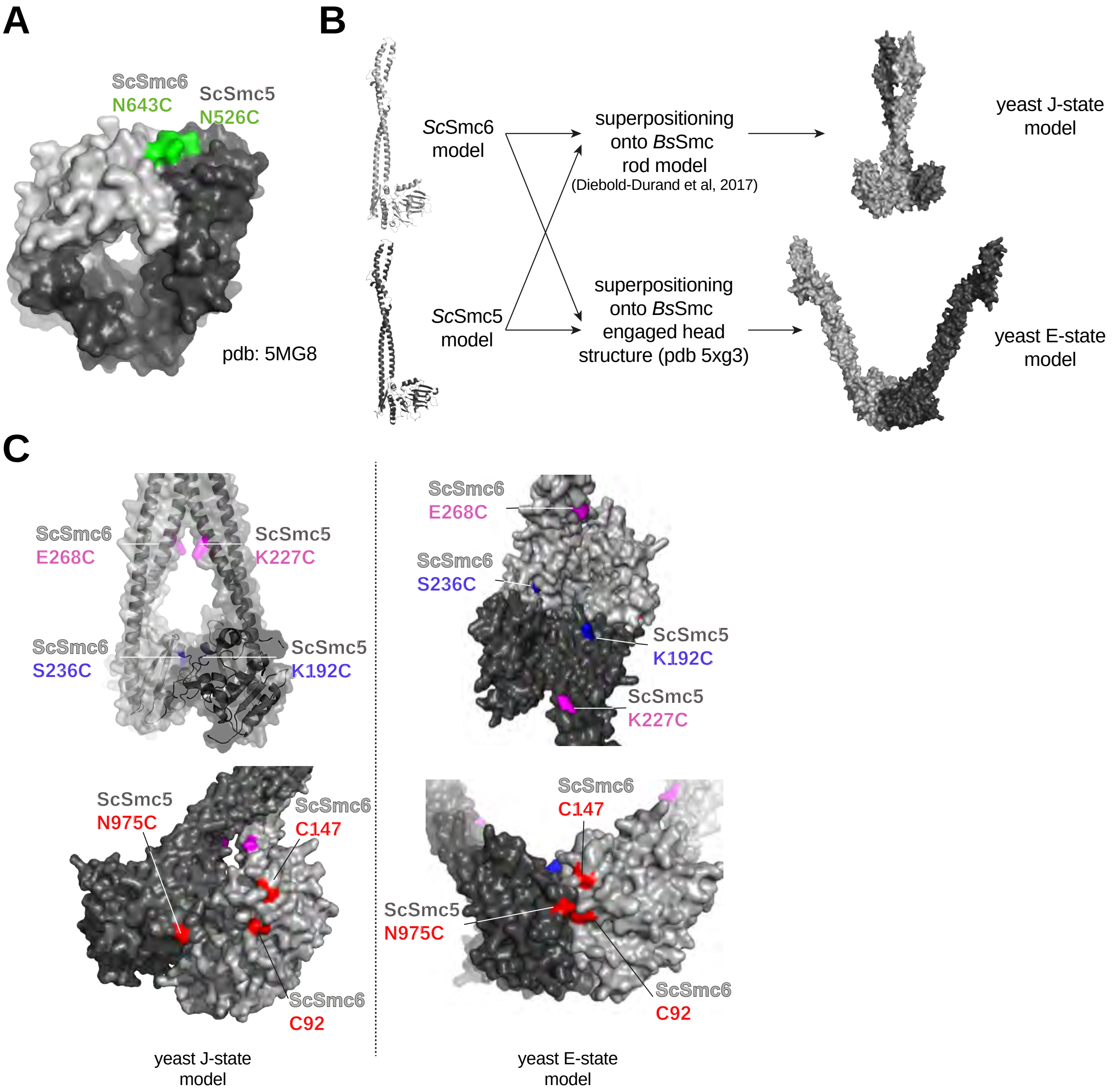


**SUPPLEMENTAL FIGURE 5**

Fig S5 Design of reporter cysteines in Smc5/6

A. Hinge-Cys residues were chosen based on a homology model of the budding yeast Smc5/6 hinge domain built from the fission yeast hinge structure (Alt et al., 2017). Residues Smc5(N526C) and Smc6(N643C) are indicated in green colours on the hinge structure in surface representation.

B. Strategy for the construction of models for the J-state and the E-state based on structural models of the archaeal Smc-ScpAB complex.

C. Positions of J-Cys (in blue colours), CC-Cys (in purple colours) and E-Cys (in red colours) residues on models of the J-state and the E-state.

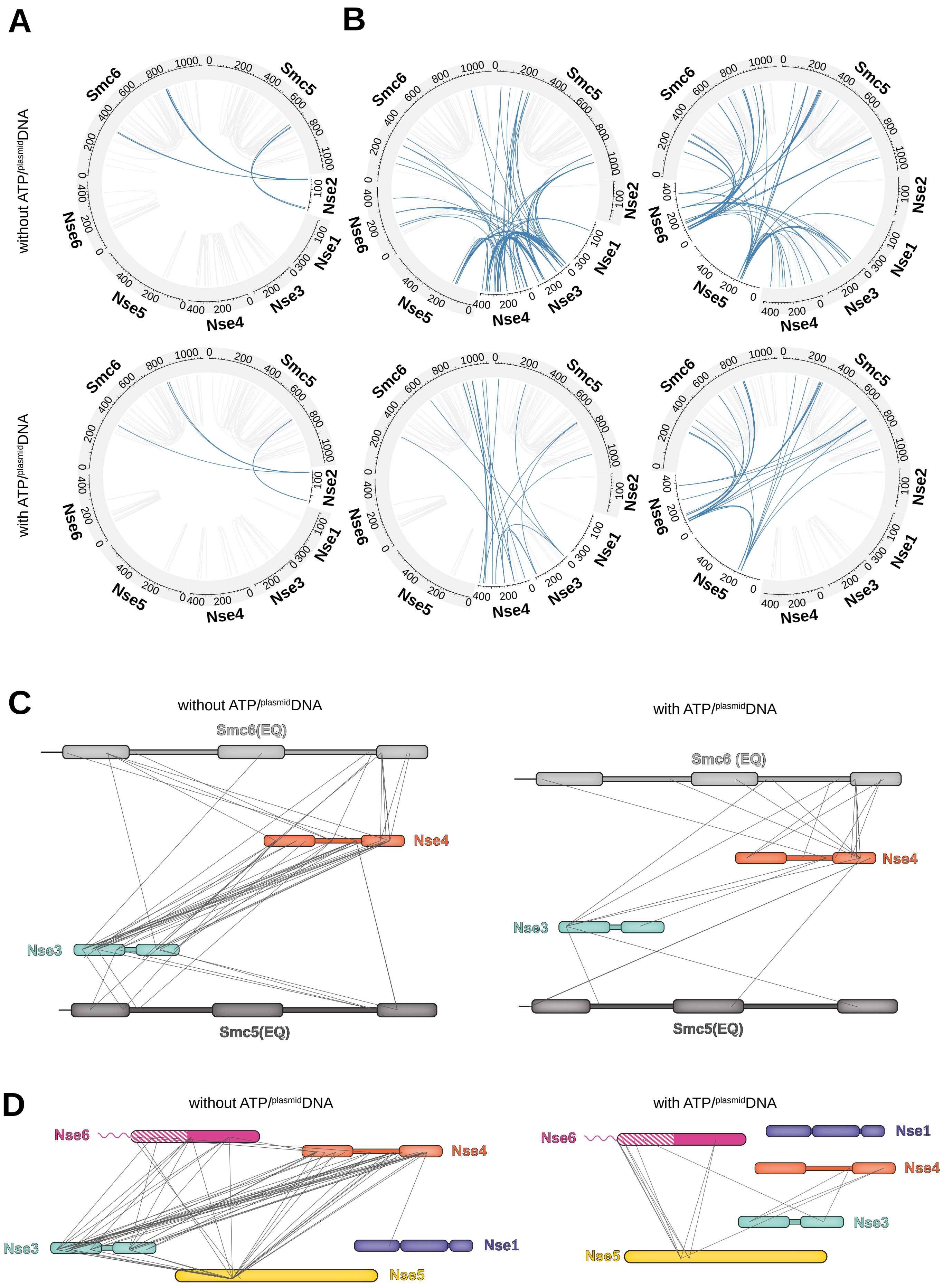


### SUPPLEMENTAL FIGURE 6

Fig S6 XL-MS of Smc5/6 octamers with and without substrates

A. Cross-links between Nse2 and Smc5/Smc6 proteins remain unaltered upon $^{plasmid}$DNA addition.

B. Changes in inter-subunit cross-links between Nse1/3/4 (left) and Nse5/6 (right) modules. Alternative representation of the same data shown in Fig 6B and 6C.

C. Cross-links between Nse3 and Nse4 and between Nse3/Nse4 and Smc5/Smc6 detected in the Smc5(EQ)/Smc6(EQ) octamer without ATP and $^{plasmid}$DNA (*left panel*) and with ATP and $^{plasmid}$DNA (*right panel*).

D. Same as in (C) for cross-links between Nse1/3/4/5/6 proteins.
